# seqsizzle: decoding complex barcode and adapter architectures in long-read sequencing data

**DOI:** 10.64898/2026.07.17.738618

**Authors:** Changqing Wang, Matthew E. Ritchie, Nadia M. Davidson

**Affiliations:** The Walter and Eliza Hall Institute of Medical Research, Parkville, Victoria, Australia; Department of Medical Biology, The University of Melbourne, Parkville, Victoria, Australia

**Keywords:** single-cell, long-read sequencing, Oxford Nanopore Technologies, Pacific BioSciences, quality control, visualization

## Abstract

Visualizing and troubleshooting long-read sequencing data is a common task when analyzing novel protocols, but identifying several primer sequences from reads exceeding 2 kb with fuzzy matching is challenging for the human eye. Here, we present `seqsizzle`, a cross-platform terminal user interface (TUI) based tool for convenient visualization of long-read sequencing data directly from the command-line, applicable to all sequencing technologies that produce `FASTQ` or `FASTA` output. `seqsizzle` allows primer sequences to be matched with user-specified fuzzy matching thresholds and highlighted with configurable colors. Additional styling can be applied to distinguish mismatches and quality scores, allowing convenient quality control of long-read sequencing data. To further aid troubleshooting, `seqsizzle` includes a built-in *k*-mer enrichment analysis, enabling the detection of unknown primer sequences or unexpected artifacts, which can then be reviewed via manual inspection. `seqsizzle` is implemented in Rust and is available open-source from https://github.com/ChangqingW/SeqSizzle.

## Background

Long-read sequencing technologies have greatly advanced our ability to profile the genome, transcriptome and epigenome, allowing gapless assemblies for creating reference genomes[1], resolving full-length transcripts at singlecell [2–4] and spatial [5] resolutions, and profiling multiple types of DNA and RNA modifications [6, 7]. Despite the technical improvements made over recent years, long-read sequencing platforms still typically produce reads with lower sequence accuracy compared to traditional short-read sequencing, with the popular Nanopore platform reported to achieve a per-base accuracy of 97-99% with its latest kits [8]. While this error rate may seem negligible at first glance, for tasks involving sequence matching, a 1% error rate could mean losing nearly 40% of reads due to sequencing errors (e.g. when matching reads that have 3 adapter sequences, each of which is 16 bases long, 1 − 0.99^3*×*16^ = 38% loss). Hence, long-read specific tools have been developed for various tasks including sequence alignment [9], single-cell barcode demultiplexing [10–12] and variant calling [13]. While these tools have been effective in their respective areas, there remains a significant gap when it comes to troubleshooting and visualizing raw long-read sequencing data quickly and interactively.

Furthermore, new sequencing protocols are rapidly being developed in various fields, such as single-cell and spatial transcriptomics, epigenomics, and clonal barcoding [14–17]. Most of these approaches incorporate multiple synthetic sequences such as primers, barcodes and unique molecular identifiers (UMIs), arranged in specific and sometimes undocumented orders. Processing sequencing data from novel protocols requires knowledge of the arrangement of these elements, for instance, assigning spatial coordinates to reads requires the processing software to identify the location of the spatial barcodes, before matching them to a coordinate dictionary. Moreover, newly developed protocols can be susceptible to higher error rates or unexpected artifacts which must be identified and removed prior to downstream analysis.

These developments create an increasing need for tools that can help troubleshoot these novel sequencing protocols, to discover artifacts and infer the read structure of the data prior to more detailed data processing steps, such as barcode demultiplexing or UMI deduplication. However, there is currently no dedicated open-source tool that can aid with these tasks, allowing visualization of raw reads directly from complex sequencing data containing errors, to offer intuitive troubleshooting of long-read protocols. In contrast, a wide range of sophisticated inspection tools exists for processed data, such as genome browsers including IGV [18]; however, these operate on trimmed and aligned data, whereas protocol troubleshooting requires inspection of unprocessed reads that will include visualization of adapters and other sequences.

To address this, we developed `seqsizzle`, an interactive terminal-based sequence reader that can detect and highlight adapter or artifact sequences with user-configurable fuzzy matching and coloring, and summarize the patterns observed in the read structure. In the remainder of this article, we describe the implementation details of `seqsizzle` and how it can be applied to perform interactive troubleshooting of a novel long-read single-cell protocol and to inspect reads from a bulk direct RNA-seq experiment.

### Implementation

`seqsizzle` provides an interactive sequence-read troubleshooting workflow, as illustrated in Figure 1A. Users can start without the adapter sequences known beforehand and use the `‘enrich’` utility to identify possible adapters, as demonstrated in Figure 1B. They can then visualize the reads with the identified primers in the main interface (Figure 1C) and configure the sequences to be highlighted (Figure 1D). Finally, the `‘summarize’` sub-command can be used to gain summarized percentages of reads in different arrangements, shown in Figure 1E.

**Fig. 1.**
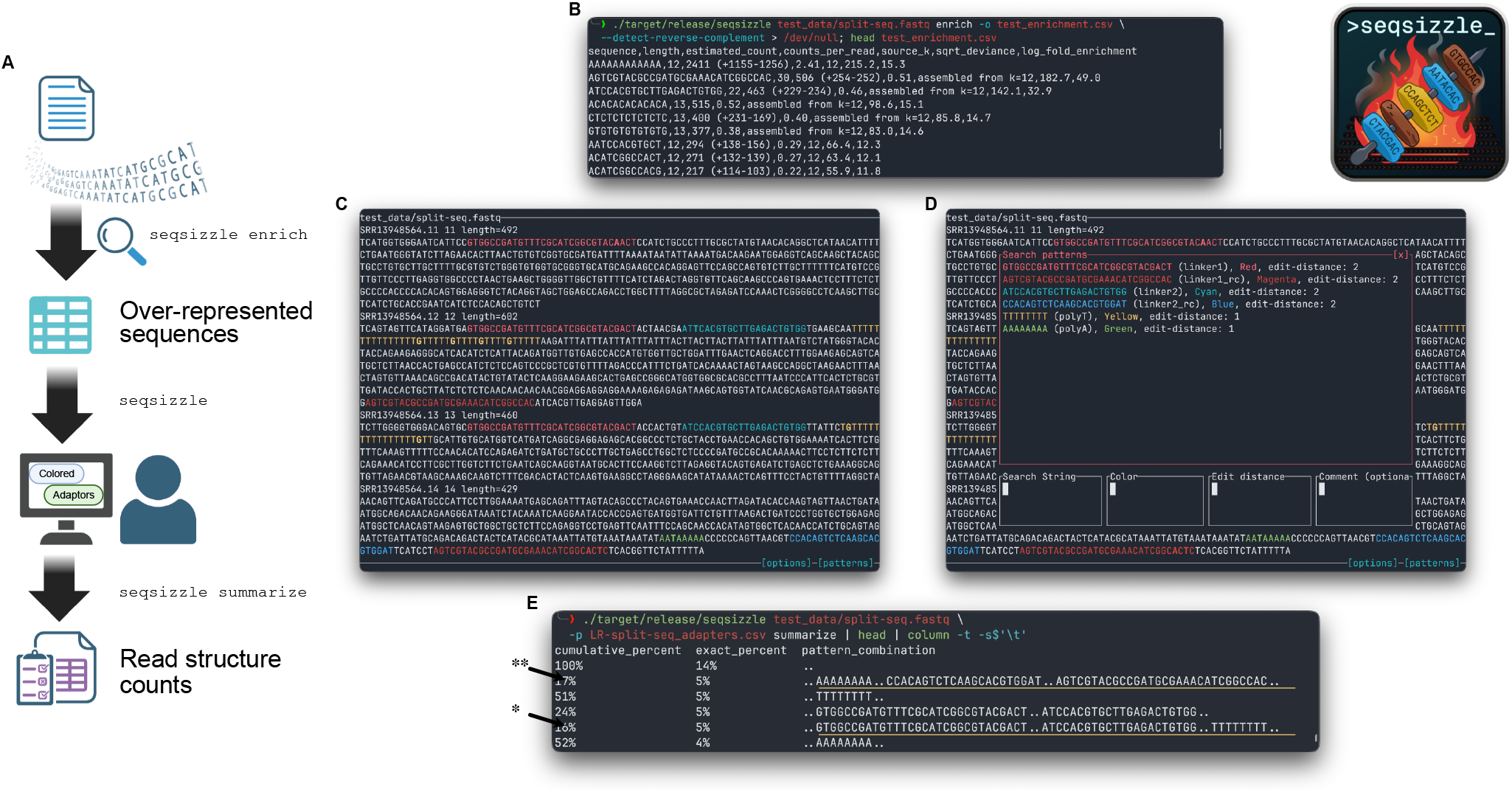
seqsizzle workflow and terminal interfaces. **A**. Flow chart of a typical `seqsizzle` workflow to visualize sequencing reads where the structure is unclear. **B**. Output of `seqsizzle`‘s `enrich` command on the first 1,000 reads from the PacBio LR-split-seq dataset. The two linker sequences and the polyA tail are reported by `seqsizzle` in the first 3 entries. **C**. Main interface of `seqsizzle`. The linker sequences, polyA and their reverse complement are highlighted in LR-split-seq reads. **D**. Search panel popup of `seqsizzle`. The LR-split-seq adapter sequences are configured here. **E**. Output of `seqsizzle`‘s `summarize` command on the first 1,000 reads from the PacBio LR-split-seq dataset. `seqsizzle` reveals that about 33% of reads are of the expected pattern: linker 1 sequence, followed by linker 2 and a stretch of polyT (the row indicated by ‘**\***’), or the reverse complement of it (indicated by ‘**\*\***’). Primer sequences were matched with the allowed edit-distance configured in **D**.

`seqsizzle` is implemented in Rust, currently supporting `FASTQ` and `FASTA` as inputs, and is distributed as a single lightweight static binary with support for Linux, macOS and Windows. Since it operates on `FASTQ` and `FASTA` directly, it is platform-agnostic and can be applied to data from any sequencing technology that outputs these formats, including Oxford Nanopore, PacBio and short-read platforms. The examples below are from PacBio and Oxford Nanopore, respectively.

Reads are matched with user-specified sequences and the allowed number of errors using Myers’ fast approximate string matching algorithm implemented in the `Rust-bio` library [19, 20]. For simplicity of implementation and ease of access from a server, the interface is implemented in the terminal, where the reads are displayed, highlighting matched regions. Mismatches are signified in bold and quality scores are stylized by italics or coloring for `FASTQ` inputs, as shown in Figure 1C.

### Using seqsizzle

The main `seqsizzle` interface displays the reads from the input file, where users may use the arrow keys to navigate through the reads and press ‘`c`‘ to toggle mouse support for clickable buttons, as shown in Figure 1C. With `FASTQ` inputs, low quality bases can be stylized to italics or alternative background coloring by toggling ‘`i`‘ (for italics) and ‘`b`‘ (for background coloring) keys. Sequences to be matched, their corresponding color and maximum errors allowed can be specified in a popup menu by pressing ‘`/`‘ (shown in Figure 1D), providing an additional `CSV` file or loading from one of the presets. To identify over-represented sequences, such as adapters and spacers, the ‘`enrich`‘ sub-command could be used, instead of launching the main interface, and a `CSV` file containing the most abundant sequences will be output.

### Sequence enrichment analysis

The `seqsizzle enrich` sub-command performs *k*-mer enrichment analysis via *k*-mer counting with increasing *k* values from a configurable arithmetic progression (8 to 12 with steps of 2 by default). For each *k* value, *k*-mers are retained only if their observed count exceeds a *Z*-score threshold based on a binomial null model assuming uniform base composition: a *k*-mer is kept when its count is at least *z* standard deviations (5 by default) more than the expected count. A fixed per-read minimum count can also be specified instead. The *k*-mers kept are then capped at the 400 most frequent per *k* by default. Additional filtering is applied to highly homopolymeric *k*-mers (when a single base accounts for ≥80% of positions in a *k*-mer), since they often accumulate large counts due to the rolling window *k*-mer counting. While only four perfectly homopolymeric *k*-mers exist per *k*, imperfect but highly homopolymeric *k*-mers can arise from sequencing errors and often crowd out the genuine enriched sequences, hence the perfectly homopolymeric *k*-mers are kept while the imperfect ones are removed.

After counting is done for all *k, k*-mers with lower *k* values are filtered if they are contained in a relatively abundant *k*-mer of larger *k* (the larger *k*-mer having counts of at least 80% of the smaller *k*-mer). Next, potential over-represented sequences longer than the maximum *k* are optionally assembled by connecting overlapping *k*-mers for the maximum *k*. Briefly, directed edges are drawn between *k*-mers with *k* − 1 bases overlap (a single-base sliding overlap) and compatible abundances (defaults to ≤20% difference in counts). The graph is traversed from unvisited *k*-mers without incoming edges, starting from the most abundant ones. The assembled sequences are extended until a branch point or cycle is encountered. The smallest count among the *k*-mers included in the assembled sequence is reported as its estimated count. Homopolymers are excluded from this step. Finally, sequences that are the reverse-complement of each other are optionally merged before the final output is written.

To accommodate protocols targeting a specific biological sequence, where the `enrich` sub-command’s report would be dominated by the targeted biological sequence, `seqsizzle` can optionally take a reference file of sequences to be excluded from *k*-mer counting. A *k*-mer exclusion set is built per *k* from the reference file, and these *k*-mers are excluded from counting during the previously described filtering stage. An error rate could also be supplied, to exclude *k*-mers within *k ×* error_rate edits from reference *k*-mers as well.

### Read summarization

The `seqsizzle summarize` sub-command takes a list of sequences to be matched with allowed errors and reduces reads to a concise description of sequence patterns, tabulating the frequency of each pattern. The supplied sequences, along with their allowed errors, are matched against the input reads using the Myers’ fast approximate string matching algorithm [19, 20]. Each read is reduced to a list of sequence matches, in the order that they occur along the read, with ‘..‘ indicating a stretch of unmatched sequence. For example, a read containing a `linker1` sequence, followed by a barcode (unmatched), and a `linker2` sequence, then another unmatched barcode, then a polyT sequence would be summarized into ‘`..[linker1]..[linker2]..[polyT]..`‘, as shown in the entry indicated by ‘**\***’ in Figure 1E. For each arrangement, the percentage of reads with that structure is counted and reported as the `cumulative_percent` and `exact_percent` columns. The `exact_percent` reports reads that fall exactly into that arrangement, with no additional matches before or after the reported arrangement. In contrast, the `cumulative_percent` also includes counts of all longer arrangements that fully contain it. For instance, additional stretches of polyT may be present after the previously mentioned arrangement, either from biological sequences or sequencing artifacts, resulting in longer arrangements.

### Example use cases

#### Recovering adapter sequences and read structure without prior information

We demonstrate how `seqsizzle` can be applied to learn about the read structure from a LR-split-seq dataset, which is a long-read single-cell RNA-seq protocol that leverages combinatorial barcoding [21]. In short, LR-split-seq involves three rounds of barcoding, with each barcode separated by a specific linker sequence. We retrieved the PacBio data from Rebboah *et al*. (SRR13948564) [21] and applied `seqsizzle`‘s `enrich` command to the first 1, 000 reads. The first 7 entries of output contain both the forward and reverse-complement sequences of the two linker sequences and the polyT tail. When supplying the `--detect-reverse-complement` option, the first 3 entries list the polyA (collapsed from both polyA and polyT sequences by the reverse-complement option) and the two linker sequences, with statistics indicating that the linker sequences both appear in about half of the reads, as shown in Figure 1B.

#### Highlighting sequences with configurable fuzzy matching and customizable color-coding

After obtaining putative adapter sequences, or when the adapter sequences are known but the user needs to inspect the reads to understand the read structure, users can launch `seqsizzle` to view the reads with the sequences highlighted, as shown in Figure 1C with the LR-split-seq data. The highlighted sequences are listed in the search panel popup shown in Figure 1D.

#### Estimating the proportion of reads with different patterns

Besides viewing the reads line by line, the `summarize` command provides a convenient pattern summary, as shown in Figure 1E, with each row of output representing a pattern or sequence combination identified in the input. For example, the 5th row (indicated by ‘**\***’) denotes reads that have a linker 1 sequence, followed by an insert of unmatched sequence, and the linker 2 sequence and then a polyT sequence, with an unmatched insert in between. Both the 5th row and its reverse complement in the 2nd row (indicated by ‘**\*\***’) are the expected arrangements in LR-split-seq, which amount to 17% + 16% = 33% of this sample of reads according to the ‘`cumulative_percent`‘ column.

Beyond finding adapters and summarizing read patterns, `seqsizzle` is useful for troubleshooting new protocols and viewing long reads in general. We demonstrate this with a Nanopore direct RNA-seq dataset on the `MOLM-13` cell line from Kim *et al*. [22], subsetting for reads from the *FLT3* gene, which is known to contain a 21-base heterozygous duplication in this cell line. The `seqsizzle enrich` Wang *et al*. | Decoding read architecture with `seqsizzle` sub-command, when excluding the *FLT3* gene reference sequence, reports the duplication site of the expected variant sequence as the top enriched sequence, as shown in Figure S1. When providing the 21-base sequence to the summarize sub-command, `seqsizzle` correctly reports that the sequence is duplicated in half of the reads, as shown in Figure S2.

#### Performance and portability

Third-generation sequencing data are commonly stored and analyzed on high-performance computing clusters rather than personal computers. To support analysis on remote devices, `seqsizzle` is implemented as a lightweight TUI program that can be run on a login node of a computing cluster without requesting additional computational resources or a graphical interface. Statically linked binaries of `seqsizzle` that are just 4.0 MB in size are provided to allow direct copying and execution on computing clusters without requiring any installation. We also tested the speed and RAM usage of the sub-command calls used in Figure 1B and 1E on an 8-core Linux instance. Over 10 runs, the enrich sub-command achieves a maximum time of 0.7 seconds and peak RAM (measured as resident set size) of 356.4 MB when processing 1,000 reads; the summarize sub-command achieves a maximum time of 54 milliseconds and peak RAM of 11.0 MB.

#### Comparison of seqsizzle and related software

A summary of comparable software is presented in Table 1. To our knowledge, `seqsizzle` is the only open-source `FASTQ` viewing software that allows multiple sequences to be fuzzy-matched and colored differently. The popular `grep` tool is very powerful at pattern matching, but is not designed for interactive visualization of sequence matches with different color-coding, nor does it support fuzzy matching. The `ugrep` [23] program provides extended support for fuzzy matching, but does not allow matching of multiple adapter sequences with unspecified order and is not designed for interactive paging through file contents. Apart from generic programs, bioinformatics-oriented tools for sequence processing and pattern searching, such as `seqkit` [24] and `cutadapt` [25] also lack interactive paging and pattern-specific colorcoding of matches. While these tools excel at processing reads, the ability to interactively inspect read structures is an important yet often overlooked feature in open-source software, prompting closed-source commercial software such as Geneious Prime [26] to fill this gap.

**Table 1.** Feature comparison of current sequence matching tools with similar functionality.

| Tool | Fuzzy multi-pattern match | Per-pattern coloring | Inter-active paging | Adapter discovery (k-mer) | Read structure summary | Open source / free | SSH friendly |
| --- | --- | --- | --- | --- | --- | --- | --- |
| `seqsizzle` | ✓ | ✓ | ✓ | ✓ | ✓ | ✓ | ✓ |
| `grep` | X | X | X | X | X | ✓ | ✓ |
| `ugrep` [23] | partial | X | X | X | X | ✓ | ✓ |
| `seqkit` [24] | ✓ | X | X | X | X | ✓ | ✓ |
| `cutadapt` [25] | ✓ | X | X | X | partial | ✓ | ✓ |
| Geneious Prime [26] | ✓ | ✓ | ✓ | partial | X | X | X |

## Discussion

`seqsizzle` addresses the need for troubleshooting of complex sequencing protocols by allowing direct inspection of the raw reads with flexible visualization of sequences, along with helper utilities to identify potential sequences of interest and provide read structure summaries. It is designed to be a lightweight and interactive troubleshooting aid, to allow data inspection for novel protocol development and help with data pre-processing.

In addition to the `FASTQ` and `FASTA` file formats that `seqsizzle` currently supports, `BAM` file input support would be a natural extension, which would allow stylizing aligned regions, so users can identify which part of their read came from their target genome, as well as visualizing alignment mismatches and base modifications. Another avenue of future development is the interoperability with BLAST or databases of common adapters and contamination. Since the `‘enrich’` command may report over-represented sequences that the user does not recognize, integration with BLAST could indicate possible sources of the sequences, saving the user from manually BLASTing and offering a more streamlined troubleshooting experience.

## Data Availability

The source code and pre-compiled binaries of `seqsizzle` are available at GitHub (https://github.com/ChangqingW/SeqSizzle), bioconda (https://bioconda.github.io/recipes/seqsizzle/README.html) and Rust’s crates.io (https://crates.io/crates/seqsizzle). The LR-split-seq data from Rebboah *et al*. was retrieved from the Sequence Read Archive (SRR13948564) [21]. The MOLM-13 cell line data from Kim *et al*. was retrieved from the Sequence Read Archive (SRR32418186) [22].

## Acknowledgments

We thank early users, including Oliver Cheng, for their feed-back and suggestions on seqsizzle. Artificial intelligence (Anthropic Claude, GitHub Copilot and Google Gemini) was used for auto-completion, code refactoring and certain feature implementations including mouse clicking support and FASTA input support, and for identifying typographical and grammatical errors in the manuscript.

## Author Contributions

C.W. developed the seqsizzle software and wrote the manuscript. M.E.R. and N.M.D. supervised the research. All authors read and approved the final manuscript.

## Conflict of interest

N.M.D. has received support from Oxford Nanopore Technologies (ONT) to present their findings at scientific conferences. However, ONT played no role in the study design, execution, analysis, or publication of this research. The other authors declare no competing interests.

## Funding

This work was supported by funding from the Chan Zuckerberg Initiative DAF, an advised fund of Silicon Valley Community Foundation (Grant No. 2019-002443 to M.E.R.), Australian National Health and Medical Research Council (NHMRC) Investigator Grants (GNT2016547 to N.M.D. and GNT2017257 to M.E.R.), the Australian Research Council (Discovery Project No. 200102903 to M.E.R.), Victorian State Government Operational Infrastructure Support and Australian Government NHMRC IRIISS. N.M.D. is also supported by the Estate of Judith Corrie Philpots.

## Notes

### Summary of Updates

Figure 1 revised; Reference URL fixed

