## supplementary information for "seqsizzle: decoding complex barcode and adapter architectures in long-read sequencing data"

### Supplementary Materials

```
╰─❯ ./target/release/seqsizzle test_data/FLT3/SRR32418186.FLT3.fastq enrich \
  --anti-reference test_data/FLT3/FLT3.fasta --anti-reference-error-rate 0.1 \
  -o SRR.csv > /dev/null; head SRR.csv
sequence,length,estimated_count,counts_per_read,source_k,sqrt_deviance,log_fold_enrichment
GAATTTGATT,10,95,0.54,10,28.4,7.6
AACAACAACAACA,13,29,0.17,assembled from k=12,20.4,11.8
TTTTTTTTTTTT,12,33,0.19,12,19.8,10.0
```

**Fig. S1. `seqsizzle enrich` recovers the expected variant sequence.** Output from `seqsizzle enrich` on Nanopore direct RNA-seq data from `MOLM-13` by Kim *et al.*, using the reference-exclusion feature. The expected heterozygous insertion sequence (first entry) is reported at around 54% frequency. Reads were filtered for the *FLT3* gene. The 21-base sequence '`TTGATTTCAGAGAATATGAAT`' is repeated in the variant allele, i.e. the variant allele has '`TTGATTTCAGAGAATATGAATTTGATTTCAGAGAATATGAAT`'. `seqsizzle`'s enrichment finds the duplicated site ('`GAATTTGATT`', as highlighted in red in the variant sequence) to be over-represented and not in the *FLT3* gene reference sequence.

```
╰─❯ ./target/release/seqsizzle test_data/FLT3/SRR32418186.FLT3.fastq \
  -p FLT3_seqs.csv summarize | column -t -s$'   '
cumulative_percent  exact_percent  pattern_combination     *
45%                 45%            ..TTGATTTCAGAGAATATGAATTTGATTTCAGAGAATATGAAT..
45%                 42%            ..TTGATTTCAGAGAATATGAAT..
100%                10%            ..
3%                  3%             ..TTGATTTCAGAGAATATGAAT..TTGATTTCAGAGAATATGAAT..
```

**Fig. S2. `seqsizzle summarize` identifies the 21-base duplication.** Output of applying `seqsizzle summarize` with the repeated 21 bases (as indicated by the box with '*'): reads with the duplication (first entry) and the sequence without duplication (second entry) each make up about half of the reads from the gene. Reads were filtered for the *FLT3* gene.
